# Input / Output Relationships for the Primary Hippocampal Circuit

**DOI:** 10.1101/2023.11.16.567451

**Authors:** BG Gunn, BS Pruess, CM Gall, G Lynch

## Abstract

The hippocampus is likely the most studied brain region but little is known about signal throughput –– the simplest, yet most essential of circuit operations –– across its multiple stages from perforant path input to CA1. Here we report that single pulse stimulation of the lateral perforant path (LPP) produces a two-part CA1 response generated by projections to CA3 („direct path‟) and the dentate gyrus („indirect path‟). The latter was by far the more potent in driving CA1 output because it engaged the massive recurrent collateral system and elicited a series of fEPSPs and spikes in CA3. The mobilization time for this stereotyped sharp wave-like response resulted in surprisingly slow throughput. The circuit did not convey high frequency LPP trains but transmitted single pulses, or bursts of pulses separated by the period of the theta wave. During these activation patterns CA1 output faithfully reproduced a version of the LPP input. We conclude that the basic hippocampal circuit, despite its considerable complexity, has a default mode in which select cortical signals are reliably transferred to output stations.

**Significance statement:** The hippocampus, a brain structure synonymous with episodic memory, is one of the most studied brain regions in neuroscience today. However, despite this intense interest, surprisingly little is known about how signals are transformed and processed by the hippocampal circuit. As a result, there are currently no “bottom up” hypotheses about how the structure supports its physiological function(s). Here, we use a novel brain slice preparation to describe the signal transformations occurring across the primary hippocampal circuit. The results identify novel circuit operations that challenge the notion of the tri-synaptic circuit and provide evidence for frequency-dependent filters that are critical for determining signal throughput. These findings provide an initial link between basic circuit function(s) and events recorded in behaving animals.

## Introduction

The signal transformations underlying the circuit level operations responsible for the critical contributions of hippocampus to episodic memory are poorly understood. Studies of rodents and humans have confirmed the expectation (from neuroanatomy) that inputs from the lateral and medial entorhinal cortices carry information about the identities and locations of cues, respectively [1–8]. Experimental work on rodents suggests that the dense recurrent collateral system in field CA3 supports encoding of the sequence in which events were sampled, a third basic element of an episode [9]. While these results identify pathways that are critical for specific memory operations, they provide little insight into the types of dynamic operations executed by the network. Data pertinent to this can be found in the large literature describing neuronal activity (spiking and local field potentials) associated with different aspects of learning (reviewed in [10]). This work has identified effects that emerge from active circuits but do not provide insights into the operational steps (i.e. signal trasformations) that generate the observed action potentials or rhythms. Thus, while much has been learned, it is not currently possible to build hypotheses of the hippocampus that begin with the fundamental signal processing (i.e., amplify, filter) that occurs within this circuit.

The present studies address a basic unresolved question about hippocampal networks: the manner in which cortical input signals are converted into reliable hippocampal output. The absence of information on this point is particularly surprising because the design of the basic circuit has been described in detail [11]. The perforant path arising from layer II of entorhinal cortex forms two branches that target the dentate gyrus (DG) and CA3, thereby initiating direct (2 synapses: cortex-CA3-CA1) and indirect (3 synapses: cortex-DG-CA3-CA1) routes to the CA1 output station [5, 12, 13]. A sparser projection from entorhinal Layer III to the most distal dendrites of CA1 adds further complexity [13]. There are detailed anatomical and physiological analyses for most of the links in these pathways [14] and numerous studies of interneurons in circuit nodes [15–20]. Despite the extensive nature of this material, it remains the case that descriptions of CA1 output produced by even very simple inputs from cortex are altogether lacking in the literature.

As a first step in addressing this issue, we devised procedures to prepare hippocampal slices that retain a sufficient population of connections between the major subdivisions to allow for analysis of throughput, from entorhinal input to CA1 output. Work of this type led to the observation that slices cut at an appropriate angle exhibit robust CA1 field potentials and spikes in response to single pulse stimulation of the lateral perforant path (LPP). However, the CA1 response was more complicated and much slower than expected from the conventional „tri-synaptic circuit‟ models of hippocampus. We accordingly conducted a series to experiments to uncover the network elements responsible for the individual components of the circuit‟s output. The results indicate that the mossy fiber (MF) projection mobilizes recurrent activity within the dense collateral system in field CA3, and this, rather than the canonical DG-CA3-CA1 circuit, drives the spiking response in CA1. Beyond this, throughput was found to be strongly dependent on the frequency, and rhythmicity, of the input signal. We conclude that the different elements of the circuit present in slices are balanced so as to allow unimpeded throughput for certain frequencies and/or patterns of cortical activity.

## Results

### Single pulse stimulation of the LPP produces a novel CA1 response

Using a novel brain slice preparation, we investigated the propagation and transformation of LPP-evoked signals across the hippocampal circuit (**Fig 1a** and **Methods**). Single-pulse LPP stimulation elicited a surprisingly complex two-part fEPSP in CA1 that was not expected given the arrangements of the two LPP inputs (i.e., direct and indirect) and the well described tri-synaptic circuit (**Fig 1a, b**). We had assumed that in a slice retaining a significant percentage of LPP inputs and the connections between subfields, CA1 responses would occur with the latency anticipated for a di– or tri-synaptic system, and display the expected summation of responses that occur in close temporal proximity (i.e., not two distinct waveforms). While the onset of the initial broad fEPSP was consistent with a di– or even tri-synaptic circuit (5.2 ± 0.3 ms; n=11 slices; **Suppl. Table 1**), the presence of a significantly delayed (∼20ms to peak) larger secondary potential was surprising and could not be accounted for by the direct (LPP-CA3-CA1) or indirect (LPP-DG-CA3-CA1) paths to CA1. Despite this, both the initial and secondary potentials were negative in the proximal apical dendrites and positive in the cell body layer, indicating that they were indeed fEPSPs generated by the dense CA3 to CA1 projections (**Fig 1b**). It thus appears that unidentified processing steps within the network produced a substantial secondary activation of CA1; a notion supported by the significantly slower slope of the delayed secondary response, in conjunction with the increased variance of this response in both dendritic and cell body layers (**Table 1; Suppl. Fig 2a, b**). Sharp waves (SPWs), spontaneous events generated in CA3 [21, 22], recorded in CA1 exhibited polarity reversals along the cell body-dendritic axis (**Fig 1c**) as expected for slices with considerable preservation of the Schaffer collateral system. There was no evidence that LPP-targeted stimulation engaged a short latency (monosynaptic) projection to CA1 from Layer III cells of the entorhinal cortex.

**Figure 1.**
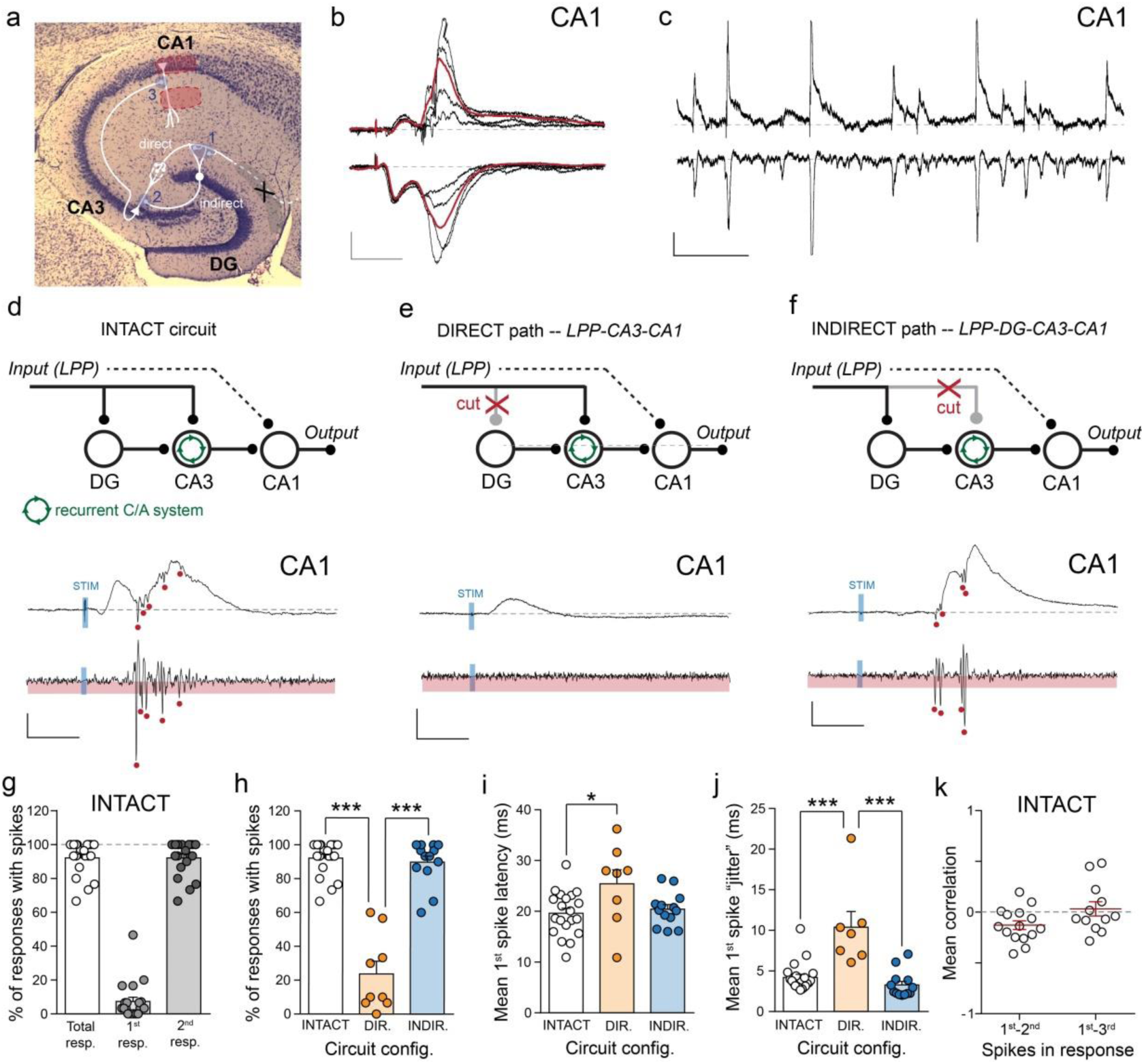
The direct and indirect LPP projections differentially engage the hippocampal circuit. **a**. Nissl stain of hippocampal slice depicting the position of stimulating electrode (X) and the two recording pipettes in field CA1 (red areas). The arrangements of the direct and indirect circuits are illustrated with the components of the tri-synaptic circuit labeled (1-3). LPP-evoked two-part fEPSPs (**b**) and spontaneous SPWs (**c**) recorded from the cell body layer (top) and apical dendrites (bottom) of CA1 (scale bars: SPW y=100μV, x=500ms; fEPSP; y=0.25mV, x=20ms). The ensemble average fEPSP is depicted in red. Schematic illustrating the circuit configuration (top) with exemplar raw (middle) and filtered (bottom,) CA1 responses for the intact circuit (**d**), direct (**e**) and indirect (**f**) paths only. Red circles depict single units within the raw and filtered signal for each (scale bars y=100μV; x=10ms). **g**. Bar graph summarizing the % of responses with spikes and their distribution across the 1^st^ and 2^nd^ components of the fEPSP. Bar graphs summarizing the % responses with spikes (**h**), latency to 1^st^ spike (**i**) and associated “jitter” (**j**) for each circuit configuration. *p<0.05; ***p<0.001 unpaired Student‟s t test. **k**. Graph summarizing the mean correlation of the 1^st^ spike amplitude with those of the 2^nd^ and 3^rd^ spikes in CA1 following LPP activation across all intact slices.

Surprisingly, evoked action potential spiking in CA1, denoting circuit output, was largely restricted to the delayed secondary component of the two-part response; this occured in 92 ± 2% of LPP-evoked responses. In contrast, spikes occurred on the first fEPSP in only 7.4 ± 3% of the time (**Fig 1d, g**). The latency to the first spike (19.6 ± 1.0 ms; n=20 slices) was much longer than anticipated for the classic tri-synaptic hippocampal circuit yet surprisingly consistent between successive pulses within and across slices (mean “jitter”: 4.2 ± 0.4 ms, n=20 slices; **Fig 1i, j**). The majority of delayed secondary responses (76 ± 5%) contained more than one spike with a mean spike frequency of 256 ± 12Hz (**Suppl. Table 2**). Two lines of evidence suggested that this high frequency spiking was generated by the near synchronous activation of a small population of CA1 pyramidal cells. First, the amplitude of the first spike was variable across trials within the same slice (CV = 37 ± 2; n=20 slices) and second, the sizes of the first and subsequent spikes were not correlated, indicating that these spikes were generated by different cells (**Fig 1k. Suppl. Fig 3a; Suppl. Table 2**).

### Origins of the two-part CA1 response to LPP stimulation

In principle, the two-part response could reflect sequential activation of the CA3 to CA1 projection by the sub-circuits initiated by the two branches of the LPP (to CA3 or DG). However, given the primary difference between the two LPP projections is the addition of a single synapse (i.e., LPP-DG) such a hypothesis does not intuitively explain the production of the two temporally distinct waveforms and the protracted delay before the onset of spiking (i.e. they are not the simply the sum of the synaptic delays). We severed specific projections using knife cuts (**Suppl. Fig 1b** and **Methods**) to test the validity of such a two pathway argument. Severing the LPP input to CA3 (LPP-CA3) or the mossy fibers (MF-CA3) did not reliably change the frequency or amplitude of SPWs recorded in CA1 (**Suppl. Fig 2c-e**), indicating that the surgical procedures only modestly disrupted the self-organizing events leading to large spontaneous depolarizations. The MF cut eliminated the second component of the CA1 response to LPP stimulation, leaving a short latency potential that corresponded to the first element of the two part response recorded in the intact slice (**Fig 1e; Suppl. Fig 2f, Suppl. Table 2**). Cutting the LPP-CA3 axons, thereby leaving intact the LPP-DG-CA3 indirect path, produced complementary results: the first component of the CA1 response was gone but the delayed secondary potential was intact (**Fig 1f. Suppl. Fig 2f, Suppl. Table 2**). The properties of fEPSPs recorded from slices lacking the indirect (i.e., MF-CA3 cut) or direct (LPP-CA3 cut) input were remarkably similar to the initial and secondary components of the two-part response of the intact circuit, respectively (**Suppl. Fig 2g-k** and **Suppl. Table 2**).

The transection experiments also provided an opportunity to test for interactions between the two pathways. Although cutting the MF-CA3 axons did produce a modest increase in the unreliable spiking response evoked by the direct path on the first of two waves of the CA1 response (**Fig 1g, h**), this reflected spikes occurring significantly later in the waveform (**Fig 1i, Suppl. Table 2**). Sectioning the LPP-CA3 axons had little if any effect on the probability, or the latency and associated “jitter”, of spikes following LPP-activation (**Fig 1h-j)**. As in the intact slice, the spike output also appeared to be driven by the near synchronous discharge from a small number of CA1 pyramidal cells (**Suppl. Fig 3b, c; Suppl. Table 2**).

Collectively, the results show that the LPP activates two intra-hippocampal pathways that sequentially engage CA1 with an unexpected delay between them. The second, delayed input drives the CA1 response and interactions between it and the direct path are surprisingly subtle.

### Variability and consistency in the CA1 response

As described, CA1 output (spikes) evoked by single-pulse LPP stimulation was largely restricted to the second, considerably delayed fEPSP. This was the case even when a sizeable number of spikes were triggered by single pulse activation of the LPP. Trial-by-trial responses did not differ greatly for individual slices (**Fig 2a, b**) but there was considerable variability between slices (**Fig 2c**). There was a significant correlation between the number of evoked spikes and the frequency of spontaneous SPWs (**Fig 2d**), which indicates that the between-slice variations were not due simply to the location of the stimulating electrode. We measured inter-spike intervals (ISIs) for those slices where LPP stimulation elicited at least three spikes in 50% of trials and found minimal differences within and between slices. The mean ISI was 4.7 ± 0.4 ms between the first and second event and 4.8 ± 0.6 ms between the second and third with these values being correlated across slices (**Fig 2e**. R^2^=0.7899). As noted, spikes within a response are generated by different neurons and it is therefore likely that the low variance spacing across trials and slices involves a yet to be identified interaction between neighboring cells. Although cutting the direct LPP projection (i.e., LPP-CA3) decreased the range of evoked spikes (**Fig 2f**), there was no evident effect on inter-spike interval for responses triggered by the LPP (**Fig 2g, h**).

**Figure 2.**
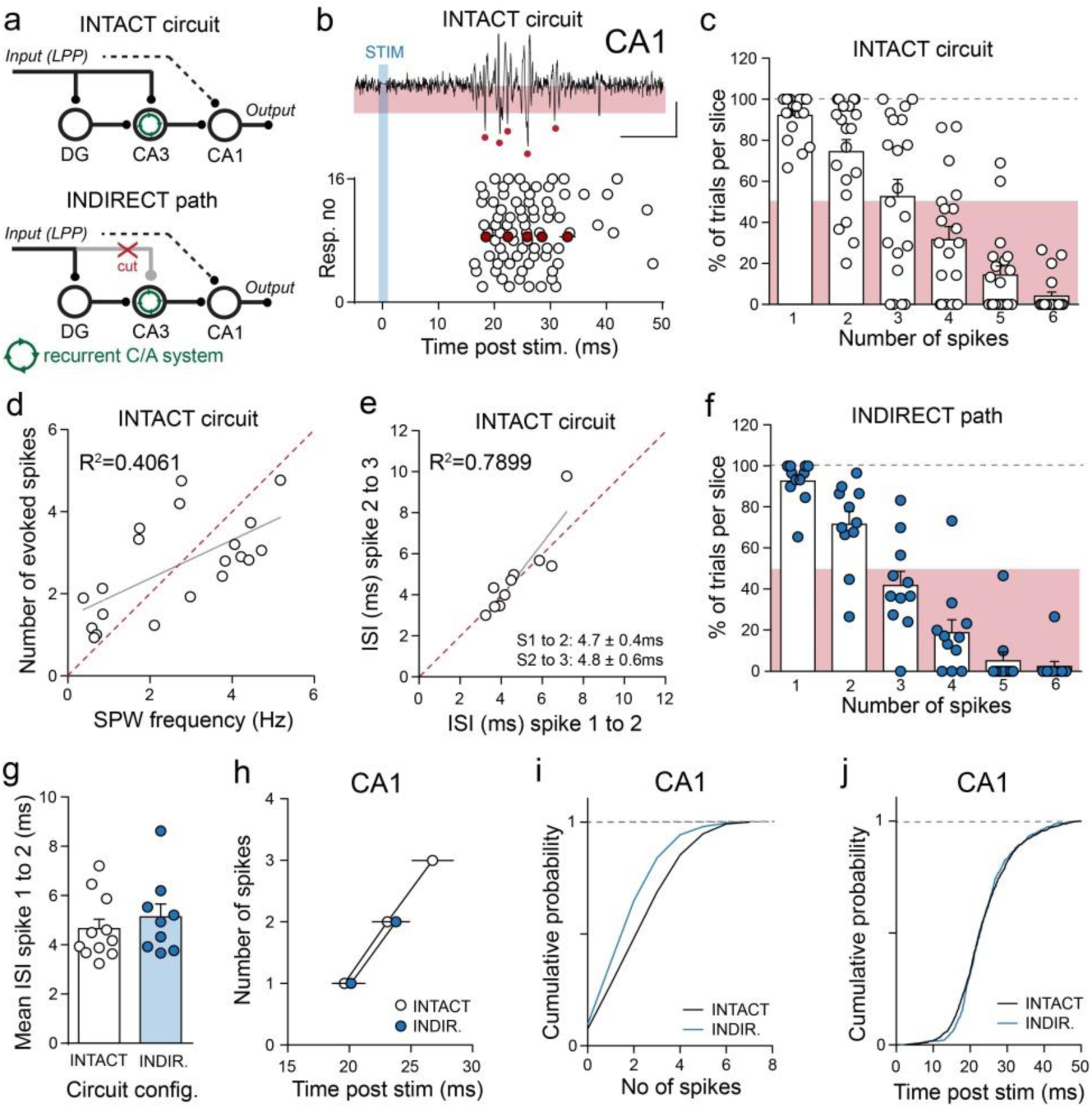
The two LPP branches interact to modestly increase the CA1 spike output to single-pulse stimulation. **a**. Schematic of the intact (top) and indirect path (bottom) circuit configurations. **b**. Example CA1 spike output from a representative intact slice (top) with the accompanying raster plot (bottom) illustrating the distribution of single units within the LPP-evoked responses (15 successive pulses: open circles). Individual spikes within the filtered trace (top) and the mean output in the raster plot (bottom) are depicted (red circles. Scale bars: y=50μV, x=10ms). Bar graph summarizing the proportion (%) of LPP-evoked CA1 responses containing 1 to 6 spikes for each slice in the intact configuration (open circle; **c**) and the indirect path only (blue circles; **f**). Bars depict the mean value (%) for each number of spikes. Scatter plots with unity line (dashed red) illustrating the relationship between the mean number of LPP-evoked CA1 spikes and the mean frequency of spontaneous SPWs (**d**) and the relationship between the intervals associated with the 1^st^ and 2^nd^ spikes and the 2^nd^ and 3^rd^ spikes (**e**). Graphs summarizing the mean ISI values between the 1^st^ and 2^nd^ spikes (**g)** and the mean number of spikes, and their temporal distribution (**h**) within the LPP-evoked CA1 response in slices containing the intact circuit (open circles) and indirect path only (blue circles). Cumulative probability plots illustrating the significant difference in the distribution of spike numbers per response between the intact circuit and indirect path (**i**; p<0.001 KS test) while no difference was observed in the temporal distribution of spikes within the waveform (**j**).

In a further test for an effect of severing the direct LPP input to CA3 on the responses generated by the indirect path (i.e., LPP-DG-CA3-CA1), we plotted the number of evoked spikes for all slices and all trials as a cumulative probability. The resultant curve for the transection group was clearly left shifted towards fewer spikes from that for the intact slices (**Fig 2i**; p<0.0001, K-S test). In contrast, curves for the onset time for evoked discharges were superimposed (**Fig 2j**). This suggests that the direct path does enhance the response to the indirect path but a large sample size is needed to detect the effect.

### The MF-CA3 link adds a complex response to the indirect path

The possibility that the indirect path, comprised of only three synapses (i.e., the„ tri-synaptic circuit‟: LPP-DG-CA3-CA1), would require almost 20 ms to drive CA1 spiking seems unlikely. This observation strongly implies that additional levels of processing within the circuit are necessary to drive throughput, and the dense recurrent commissural-associational (C/A) system in field CA3 provides an attractive candidate for performing such operations. Indeed, prior studies have demonstrated that MF stimulation is capable of driving a secondary recurrent fEPSP [23], although little is known about the interaction(s) between these two responses and their respective contribution to hippocampal throughput. First, we stimulated the MFs deep within the DG hilus and recorded from the cell bodies and proximal apical dendrites of field CA3b (**Fig 3a**). Consistent with prior studies [23] single-pulse stimulation produced a small, short latency mono-synaptic fEPSP, which was followed by a large, long duration fEPSP that reversed in the proximal apical dendrites, indicating the presence of a large current sink at this site (**Fig 3b**). The peak of the after-potential, whether recorded from the pyramidal cell layer or str. radiatum, was delayed by ∼4ms from the mono-synaptic MF response (**Fig 3c, Suppl. Table 3**). This interval combined with the presence of soma-dendrite dipole indicates that the secondary response was generated by the CA3 C/A projections (MF-CA3-CA3).

**Figure 3.**
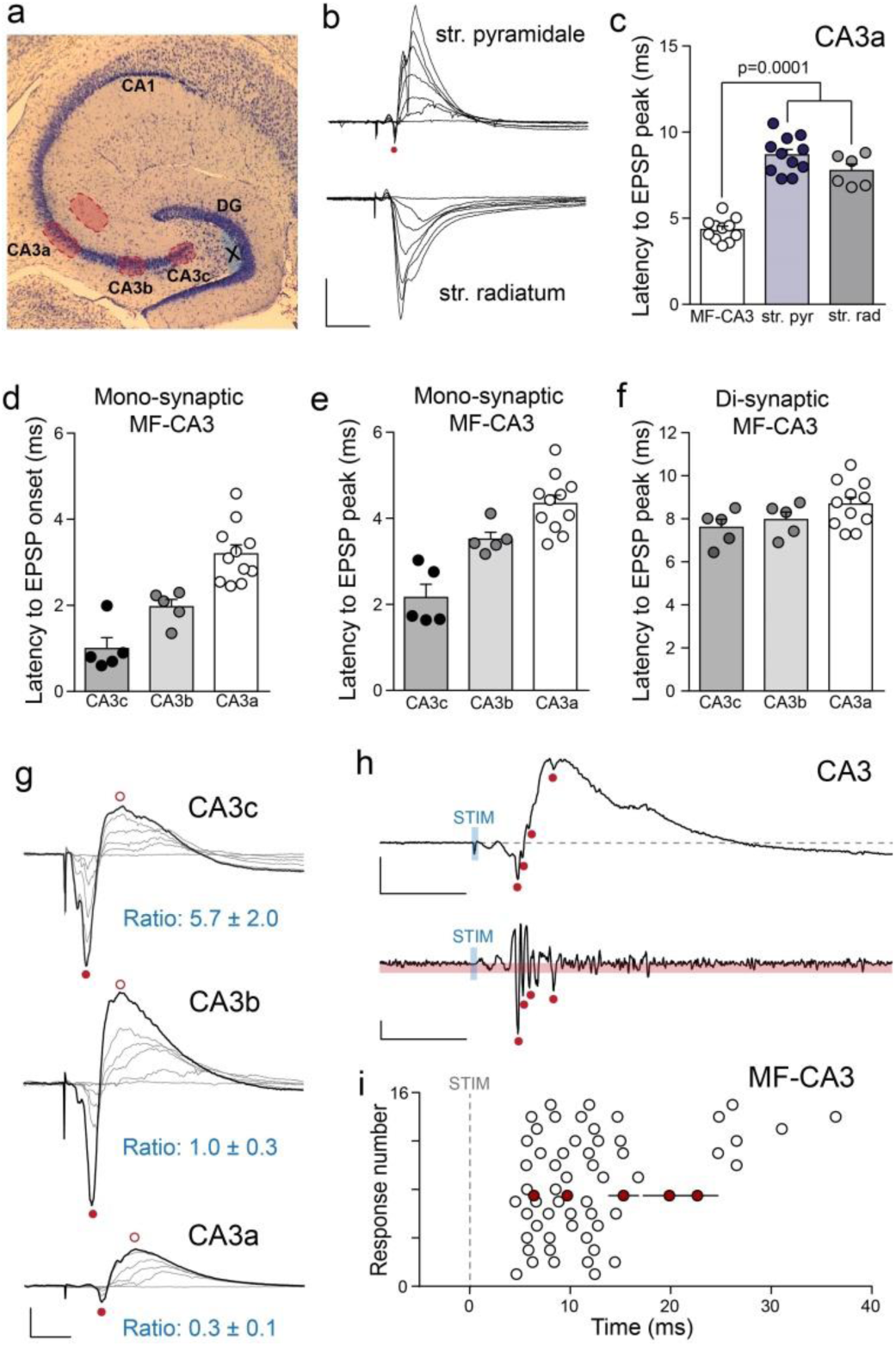
The mossy fiber synapse reliably engages the CA3 recurrent system. **a**. Nissl stain of hippocampal slice depicting the position of the stimulating electrode proximal to the granule cell layer and the typical locations of the two recording pipettes in str. radiatum and the PC layer of CA3a. The positions of recording pipettes across the proximo-distal axis of CA3 (i.e., CA3c, CA3b) are also illustrated. **b**. Representative MF-evoked fEPSPs recorded from the PC layer (top) and apical dendrites of CA3a across a range of stimulation intensities. The short latency, mono-synaptic MF-evoked fEPSP is depicted (red circle) in the PC layer (y=2mV, x=10ms) **c**. Bar graph summarizing the latency to fEPSP peak for the mono– and di-synaptic MF-evoked responses. Bar graphs summarizing the latency to onset (**d**) and peak (**e**) of the mono-synaptic MF-CA3 response as well as the latency to peak of the di-synaptic fEPSP (**f**) across the CA3 proximo-distal axis: CA3c, CA3b and CA3a. **g**. Representative MF-evoked response recorded from CA3c (top), CA3b (middle) and CA3a (bottom), with responses to increasing stimulation intensities depicted (grey traces). The peak mono– and di-synaptic responses used to calculate the ratio are depicted with closed and open red circles respectively (scale bars: y=2mV, x=10ms). **h**. Exemplar raw (top) and filtered (bottom) MF-evoked CA3 response, with single units depicted (red circles, Scale bars: y=1mV and 50μV, x=10ms). **i**. Raster plot from a representative slice illustrating the distribution of single units within the MF-evoked CA3 responses across 15 successive pulses (open circles) and with the mean slice output depicted (red circles).

The nature of MF projections, being thin and poorly myelinated, suggests signal conduction along the proximo-distal axis of CA3 will be slow. This is relevant to the question of the long delay in the indirect path because the sampled CA1 region (CA1c) receives most of its input from the more distal regions of CA3 (i.e., CA3a). To address this, we measured the amplitudes and onset times for the monosynaptic MF response and the subsequent C/A potential at three proximo-distal points in CA3 (i.e., CA3c-CA3b-CA3a). As expected, the MF response was considerably delayed in the distal aspect of CA3 relative to values recorded proximal to the DG (p=0.0001, one–way ANOVA; **Fig 3d, e**). In contrast, a tendency towards greater delay with distance for the peak of the secondary potential did not approach statistical significance (P=0.1104) **Fig. 3f**). The amplitude of the MF response underwent a marked decrease from the middle to distal end of CA3 (**Suppl. Table 3**) presumably due to i) a decreasing percentage of stimulated fibers that remain in the plane of the slice between the stimulation and recording sites and ii) the decline in basal MF terminals along the CA3 proximo-distal axis [24, 25]. The results for the C/A fEPSP did not parallel those for the MFs. Peak amplitude was greatest at the mid-point of the CA3 axis (CA3b) and not nearly so different between the distal and proximal ends as was the case for the MF potential (**Fig 3g**). The recurrent collateral system is comprised of local and long range fibers arising from sites throughout the field [26–28]. The gradient of secondary potentials just described could reflect a forward bias (towards CA1 rather than the DG) in the long collateral branches.

Spiking initiated by stimulation typically began with the MF response and continued through the rising phase and up to the peak of the secondary potential (**Fig 3h**). A trial-by-trial record for a typical slice showed that pyramidal cell discharges typically continue for about 10ms (**Fig 3i**). This block of time, when added to delays required by processing in, and the subsequent transmission through the LPP-DG-MF pathway, may provide a plausible explanation as to why almost 20 ms is required for the indirect path to engage CA1. We therefore propose that CA1 is not directly triggered by the monosynaptic response of CA3 to MF input (i.e., the tri-synaptic circuit), but rather by local cycling of the associational CA3-CA3 connections set in motion by the MFs. These features combined with the conduction velocity of the MFs account for the slow throughput that is a singular feature of the hippocampal circuit.

### The LPP triggers a prolonged but stereotyped CA3 response

We compared LPP-driven activation of CA3 with that obtained following direct MF stimulation (above). Single LPP pulses produced a striking response. Recordings in the CA3 cell body layer were characterized by an initial large potential followed by a series of smaller fEPSPs most of which were accompanied by an action potential (**Fig 4a**, **Suppl. Table 4**). Indeed, the LPP pulse triggered more spikes than did direct MF stimulation, as can be seen in trial-by-trial measures for representative cases (**Fig 4b**) and group summaries (**Fig 4c, d**). Note that although the stimulation triggered more CA3 action potentials than it did in CA1 (see **Fig 1h**), this spiking appeared similarly to be generated by a small population of CA3 pyramidal cells (**Suppl. Fig 4a, b**). As expected, the onset of the response to LPP stimulation was delayed and more variable than that for the direct MF activation (**Fig 4e, f)**.

**Figure 4.**
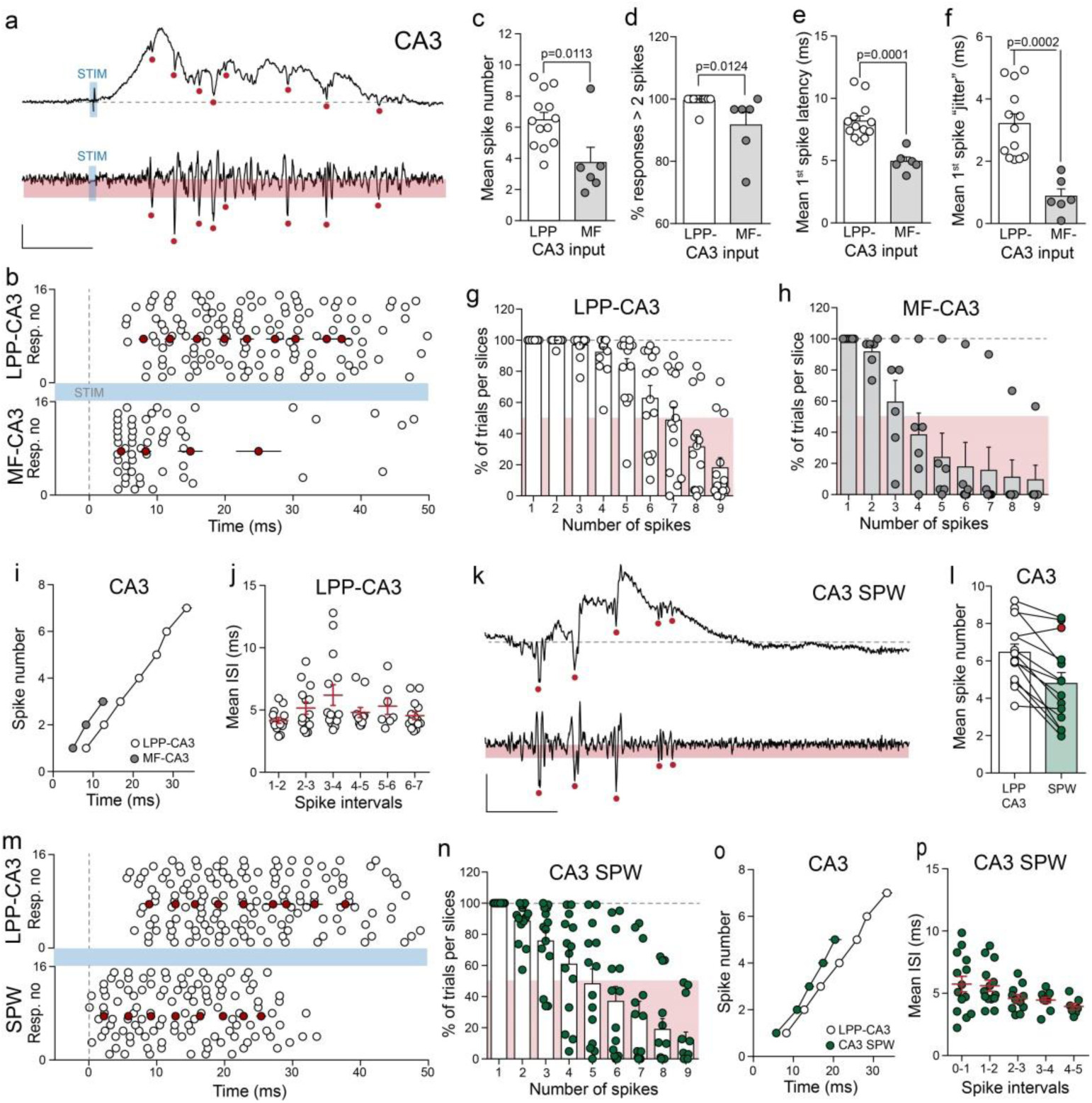
LPP activation produces a prolonged spike output from CA3 that is highly stereotyped in nature. **a**. Exemplar raw (top) and filtered (bottom) LPP-evoked response recorded from CA3a. The single units are depicted (red circles) for each (scale bars: y=200μV and 50μV, x=10ms). **b**. Raster plots illustrating the distribution of single units within the LPP-evoked (top) and MF-evoked (bottom) CA3 responses (15 successive pulses: open circles) derived from representative slices. The mean output for each slice is depicted (red circles). Bar graphs summarizing the mean number of spikes per response (**c**), the proportion of responses with >2 spikes (**d**) as well as the mean 1^st^ spike latency (**e**) and associated jitter (**f**) for LPP (open circles) and MF-evoked (grey circles) CA3 responses. Bar graphs summarizing the proportion (%) of CA3 responses containing 1 to 9 spikes for each slice following LPP (**g**; open circles) and MF (**h**; grey circles) stimulation. The bars depict the mean value (%) for each number of spikes. **i**. Graph summarizing the mean number of spikes, and their temporal distribution within the LPP-evoked (open circles) and MF-evoked (grey circles) CA3 response. **j**. Graph summarizing the mean inter-spike intervals (ISI) between successive spikes (spikes 1 to 7) across the LPP-evoked waveform for each slice. Red bars depict the mean values. **k**. Exemplar raw (top) and filtered (bottom) spontaneous SPW recorded from CA3a. The single units are depicted (red circles) for each (scale bars: y=100μV, x=10ms). **l**. Graph of the within slice comparison between the mean number of LPP-evoked spikes (open circles) with those associated with SPWs (green circles). Red circle depicts the case where SPW spiking was elevated. **m**. Raster plots illustrating the distribution of single units within the waveform of LPP-evoked CA3 responses (top) and SPWs (bottom; 15 successive responses for each: open circles) derived from representative slices. The mean output for each slice is depicted (red circles). **n**. Bar graph summarizing the proportion (%) of CA3 SPWs containing 1 to 9 spikes for each slice, with bars depicting the mean value (%) for each number of spikes. Note the similarity in the distribution *cf* LPP-evoked response (**g**). **o**. Graph summarizing the mean number of spikes, and their temporal distribution within the waveform of LPP-evoked responses (open circles) and spontaneous SPWs (green circles) recorded from CA3a. **p**. Graph summarizing the mean inter-spike intervals (ISI) between successive spikes (spikes 1 to 5) associated with SPWs for each slice. Red bars depict the mean values.

The CA3 spike output in response to LPP stimulation exhibited the same low within slice variance evident in CA1 (**Fig 4b**). As anticipated given the lower number of MF-evoked spikes, the range of spikes was decreased compared to those elicited by LPP stimulation (**Fig 4g, h**). Indeed, following LPP activation at least 50% of trials elicited 7 spikes or more in at least half of the slices tested, while MF stimulation only evoked 3 spikes (**Fig 4g-i**). Despite these differences, the ISI between spikes was remarkably similar in responses evoked by the two inputs (**Fig 4i**). The mean ISI for LPP stimulation was 5.0 ± 0.3 ms and this was surprisingly consistent across successive inter-spike intervals (**Fig 4j; Suppl. Fig 4c**). The intervals between the 1^st^ and 2^nd^ spikes were modestly, yet significantly different (LPP: 4.5 ± 0.3 ms, MF: 3.3 ± 0.3ms; p=0.0228 unpaired t test), whereas no differences were evident for the interval between the 2^nd^ and 3^rd^ spike (LPP: 4.2 ± 0.2 ms, MF: 5.2 ± 1.1 ms; p=0.1606 unpaired t test). These findings suggest that the recurrent C/A system may be predisposed to self-organization and subsequent generation of stereotyped responses. If so, then it is possible that the large spontaneous SPWs in CA3 would display a similarly stereotyped spiking profile. We tested this using the same filtering procedures employed in the stimulation experiments and estimating the start of the SPW from the field potential trace **(Fig 4k**). Within-slice comparisons confirmed that fewer spikes were present during SPWs than were elicited by LPP stimulation (**Fig 4l, m**; p=0.0002 paired t test). As with the LPP-evoked responses, the variability in spikes associated with SPWs was low within each slice (**Fig 4m**), but considerably larger across slices (**Fig 4n**). In general, the ISI values for spikes within a SPW were remarkably similar to those obtained following LPP stimulation (**Fig 4o**). However, in contrast to LPP-evoked spikes the variance in the ISIs between successive spikes (i.e., 1^st^, 2^nd^, 3^rd^) associated with SPWs actually decreased, with a tendency for the mean ISIs to be slightly shorter than for those observed following LPP stimulation (**Fig 4j, p**).

### Throughput from the LPP to field CA1 is frequency-dependent

We next investigated the output of the circuit in response to LPP stimulation frequencies that are both prominent during behavior and widely utilized in models of cortical and hippocampal computations (refs). Activation at the theta frequency (5Hz) produced spiking by CA1 pyramidal cells that was maintained throughout a train of ten pulses with or without the direct input to CA3 (**Fig 5a, b**). Results for all slices in both groups confirmed that throughput (i.e., CA1 cell firing) was almost always present for each pulse in the train (**Fig 5b, Suppl. Fig 5a)**, with the number and patterning of spikes elicited by the 10^th^ pulse not being reliably different than those for the 1^st^ pulse (**Fig 5c, d, Suppl. Fig 5b**). The similarity between the first and last responses extended to the onset time and its associated variance for the spiking response, which was virtually identical for the 1^st^ and 10^th^ pulses in the intact slices (**Fig 5e left, Suppl. Fig 5e**). Interestingly, transection of the direct LPP input shortened the delay to 10^th^ pulse relative to the 1^st^ (**Fig 5e, right, Suppl. Fig 5c**). The number of action potentials in the 50ms following the first and last pulses correlated across slices with the responses collected during baseline recording (**Fig 5f**), indicating again that repetitive stimulation at 5Hz does not significantly change the resting properties of the system.

**Figure 5.**
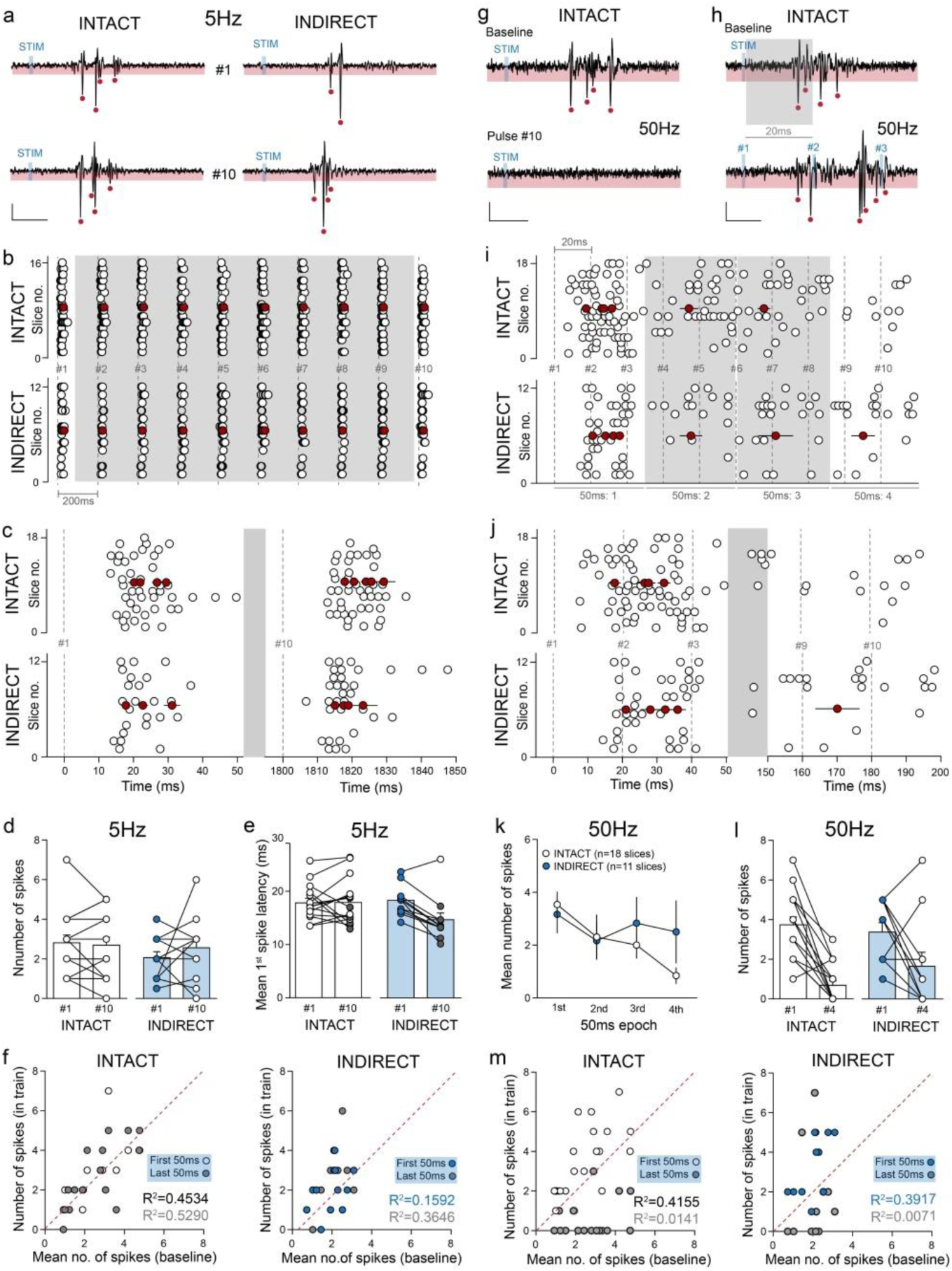
The hippocampal circuit operates as a low pass filter in response to repetitive stimulation. **a**. Representative filtered LPP-evoked CA1 responses for the 1^st^ and 10^th^ pulse of a 5Hz (10 pulse) stimulation train in the intact circuit (left) and the indirect path only (right; scale bars: y=50μV; x=10ms). **b**. Raster plot of LPP-evoked CA1 spiking across the 5Hz stimulation train for each slice (open circles) recorded with the intact circuit (top) and indirect path only (bottom). The distribution of CA1 spiking during the 1^st^ and 10^th^ response for each slice (both circuit configurations) is illustrated on an expanded time scale (**c**). Red circles depict the mean spike distribution for all slices in both. Bar graphs of the mean number of CA1 spikes (**d**) and the mean 1^st^ spike latency (**e**) on the 1^st^ and 10^th^ stimulation pulses of a 5Hz train in the intact circuit (white) and with the indirect path only (blue). **f**. Scatter plots with unity line (dashed red) summarizes the mean number of CA1 spikes evoked during the single-pulse baseline period and those during the 1^st^ and 10^th^ pulse of a 5Hz stimulation train for each slice in the intact circuit (left) and with indirect path only (right). **g.** Representative filtered LPP-evoked CA1 baseline response (top) and 10^th^ pulse (bottom) of a 50Hz (10 pulse) stimulation train in the intact circuit. **h**. Representative filtered LPP-evoked CA1 baseline response (top) and the response over the first 50ms (3 pulses) of a 50Hz (10 pulse) stimulation train in the intact circuit (scale bars both: y=50μV; x=10ms). **i**. Raster plot of LPP-evoked CA1 spiking across the 50Hz stimulation train for each slice (open circles) recorded with the intact circuit (top) and indirect path only (bottom). The distribution of CA1 spiking during the 1^st^ and last 50ms periods of the stimulation train for each slice (both circuit configurations) is illustrated on an expanded time scale (**j**). Red circles depict the mean spike distribution for all slices in both. **k**. Graph summarizing the number of evoked CA1 spikes in each 50ms epoch across the 50Hz stimulation train in the intact slice (open circles) and indirect path only (blue circles). **l**. The mean number of spikes evoked in the 1^st^ and last 50ms of the 50Hz train for the intact circuit (left) and indirect path only (right). **m**. Scatter plots with unity line (dashed red) summarizes the mean number of CA1 spikes evoked during the single-pulse baseline period and those during the 1^st^ and last 50ms epochs of a 50Hz stimulation train for each slice in the intact circuit (left) and with indirect path only (right).

Repeating the experiments with LPP stimulation at 50Hz (gamma) yielded a startling result: there was typically no spike response at the 10^th^ pulse in the train (**Fig 5g**, bottom). Combined with the theta results this outcome indicates that the hippocampal circuit operates as a potent low pass filter (i.e., 5 Hz activity drove continued throughput to CA1 whereas 50 Hz activity did not). Representative traces showed that the filtering was not present on the response to the 2^nd^ pulse (**Fig 5h**, bottom), whereas pulse by pulse analyses for all slices in the group indicated that suppression emerged by pulse #3 (**Fig 5i, Suppl. Fig d**). While qualitatively, the effects of transection appeared minimal (**Fig 5i, Suppl. Fig e, f**), efforts to quantitatively assess possible effects were complicated by the similarity between the latency to spike (19.6 ± 1.0 ms) and the 20 ms intervals in a 50Hz train. To circumvent this problem, we divided the 200ms gamma train into 50ms epochs, which was the length of the sampling period used to assess CA1 spike output to baseline and 5Hz stimulation (and **Suppl. Methods**). The results showed that there was a ∼85% reduction in the number of evoked spikes between the first and fourth stimulation blocks (**Fig 5k, l**). The number of evoked spikes in the first 50ms epoch was tightly correlated with, and greater than, the number elicited by single pulses during baseline stimulation. The last block of responses was predictably reduced from the pre-train values (**Fig 5m left**). A similar pattern held for slices in which the direct path had been severed (**Fig 5m right**). Note that, consistent with single-pulse stimulation, activation of the direct LPP input alone (i.e., MF cut) did not reliably drive CA1 spikes at either of the two frequencies (**Suppl. Fig 5a-d, f**).

### Theta-gamma LPP stimulation obviates low pass filtering by the circuit

It is puzzling that the cognitively critical gamma rhythm is not relayed across the multiple stages of the hippocampal circuit. Possibly relevant to this, gamma activity often occurs in brief bursts separated by the period of the theta wave [29] and, as described (e.g., **Fig 5h**), two pulses applied to the LPP at 50Hz elicit a robust response in CA1. The question then arises as to whether a series of short bursts produces reliable throughput as opposed to engaging potent filtering mechanisms. We tested this by applying three pulses to the LPP at 50Hz and then repeating this at 200ms intervals for a total of five gamma bursts. LPP activation with such theta-gamma stimulation produced reliable CA1 spike output in response to each individual burst with the response to the 5^th^ burst being equal to or greater than that to the first (**Fig 6a**). A burst-by-burst analysis across all slices confirmed that each set of three pulses elicited a robust response in CA1 (**Fig 6b**). These data also suggested that while removing the direct path reduced the amount of spiking to a given burst, there was an increase in bursting across successive bursts (**Fig 6b, c**). An analysis of individual burst responses strengthened elements of this impression. While the difference in the mean number of spikes across the 50ms period following initiation of each gamma burst approached significance (intact: 3.8 ± 0.1; indirect: 3.1 ± 0.3, p=0.0680 unpaired t test), the absolute change in spike number across successive bursts differed significantly between the two groups (**Fig 6d**, F_4,100_=2.478, p=0.0488 two-way ANOVA). These results constitute a clear instance in which the direct path influences an operation of the indirect path.

**Figure 6.**
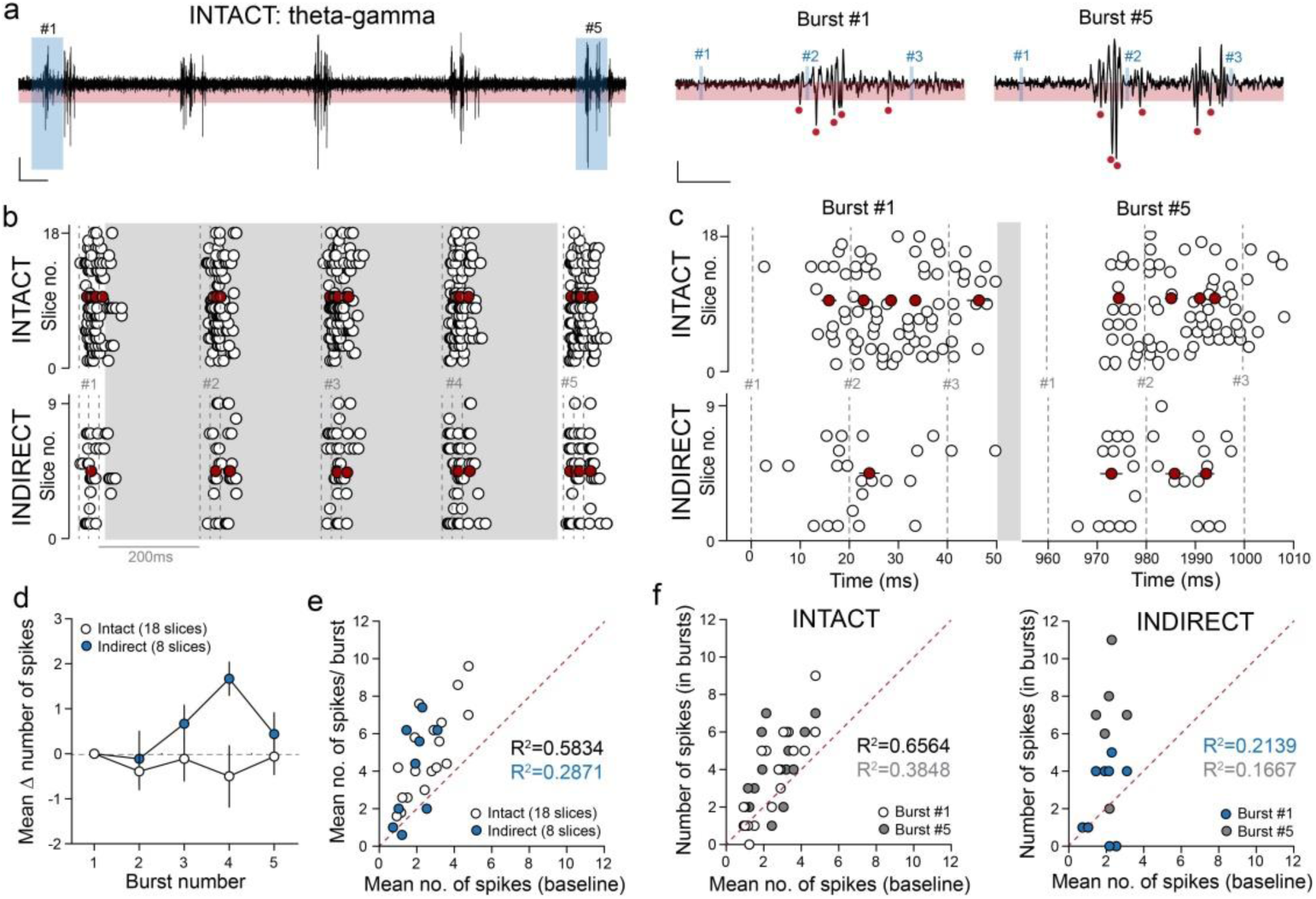
Theta-gamma stimulation obviates the low pass filter. **a**. Representative filtered CA1 response to theta-gamma (5 bursts) stimulation of the LPP input (left) in an intact slice preparation. The responses to burst #1 and #5 are illustrated on an expanded time scale (right; scale bars: y=50μV, x=100ms and 10ms). b. Raster plots of LPP-evoked CA1 spiking across the theta-gamma stimulation train for each slice (open circles) recorded with the intact circuit (top) and indirect path only (bottom). The distribution of CA1 spiking during the 1^st^ and 5^th^ burst for each slice (both circuit configurations) is illustrated on an expanded time scale (c). Red circles depict the mean spike distribution for all slices in both. d. Graph summarizing the mean change in the number of evoked spikes for each burst recorded from intact slices (open circles) and those containing only the indirect path (blue circles). Note the lack of change in the intact slice, and significant increase across bursts in the absence of the direct input. Scatter plots with unity lines (dashed red) summarizing the mean number of CA1 spikes evoked during the single-pulse baseline period with (e) the mean number of spikes per burst in the intact circuit (open circles) and with indirect path only (blue circles) as well as with (f) the mean number of CA1 spikes during the 1^st^ and 5^th^ burst of a theta-gamma train for each slice in the intact circuit (left) and with indirect path only (right). Note the significant increase in spike number associated with each burst.

As anticipated, the number of evoked spikes was extremely stable over the course of five theta-gamma bursts in the intact slices; a feature that was less evident in slices lacking the direct LPP input (**Fig 6d; Suppl. Fig. 5g, h**). Notably, between slice variations in spike responses were correlated with the magnitude of the responses to single pulse stimulation during baseline recording (**Fig 6e, f**). As with single pulse stimulation, 5Hz and 50Hz LPP stimulation to the direct input alone was ineffective at driving CA1 spiking (**Suppl. Fig 5i**). We conclude from these results that theta-gamma signals from the lateral entorhinal cortex are reliably transferred across the circuit with minimal distortion and that the direct path promotes this indirect path function. Indeed, the strongest influence of the direct path on throughput of LPP signals emerges during theta-gamma input.

## Discussion

A goal of this project was to obtain a first description of how simple inputs are transformed across the multiple stages of the primary hippocampal circuit. It is expected that such information will be of considerable utility in model building and the related development of bottom-up hypotheses about hippocampal functions. The results were unexpected and largely inconsistent with usual assumptions about the operation of intra-hippocampal networks. Single-pulse stimulation of the LPP, one of two primary inputs from entorhinal cortex, evoked two distinct fEPSPs in CA1 that were separated by a surprisingly long delay. Cell firing, the marker for CA1 output, occurred almost exclusively on the second waveform in the two-part CA1 response, and displayed surprising consistency in terms of the latency to onset and length of intervals between events. Sectioning studies revealed that the LPP branch projecting directly to CA3 (i.e., LPP-CA3 axons) initiated the first potential whereas the second waveform along with its associated spiking was driven by the LPP-DG-CA3 sub-circuit. However, these experiments did not of themselves explain why the inclusion of an additional synapse (i.e., DG-CA3) produces uncommonly slow yet temporally consistent throughput.

The absence of a correlation between spike amplitudes led to the conclusion that the CA1 response was the product of different neurons emitting single spikes. The net result was a high frequency output signal from a local population of cells. This pattern undoubtedly reflects interactions between dendritic properties, local circuit operations, and the spatio-temporal organization of input from CA3. Of interest in this regard, pyramidal cells are commonly described as integrators, that can process temporally dispersed inputs, or as coincidence detectors that preferentially respond to near synchronous signals [30, 31]. Which of these two functional modes the cells display will strongly influence the type of input needed to trigger spikes, and the degree of synchrony between the input and target cell output. Although integration of dendritic inputs is complex [32, 33], the difference between an integrator and coincidence detector can be viewed simplistically as the presence in the latter of a slow outward (hyperpolarizing) current at peri-threshold potentials [31]. Typically these currents are mediated by dendritic voltage– and Ca^2+^-dependent K^+^ channels that are activated by sub-threshold depolarization evident during high levels of stochastic synaptic input [34]. Given the considerable spontaneous activity in CA3, it can be assumed that these pyramidal cells receive a significant degree of continuous excitatory input and accordingly operate with coincidence detector traits. Such a scenario provides a plausible explanation for the low variability in the latency to CA1 spiking as well as the observation that the output is mediated by individual cells emitting a single spike [31].

The above argument suggests that activation of CA1 pyramidal cells requires afferent input that arrives in close temporal proximity. In accord with this, single-pulse LPP stimulation produced a spike output from CA3a (the subfield innervating the CA1 segment used for output measurements) lasting for ∼20 ms at a frequency in excess of 200Hz. The response coincided with a slow wave with a dipole corresponding to that generated by direct activation of the recurrent collateral system. We conclude that the relay of signals from cortex through CA3 is not mediated by the direct response to the mossy fibers but instead by a rapid build-up of local spiking driven by feedback (recurrent) excitation. The amplification step provided by this system offsets the step-down action associated with the sparse firing of DG granule cells [35, 36] and thereby enables signal throughput. The time required for recurrent feedback does however result in the unexpectedly slow throughput across the circuit.

The CA3 response to LPP stimulation was also unusual in that there was very little variability within and between slices in the intervals separating action potentials, which as in CA1 were generated by different cells. Moreover, the direct stimulation of the mossy fibers produced the same ISI as found with LPP activation and a similar value was recorded for spiking during spontaneous SPWs. The stereotyped response of CA3 to any form of activation could reflect the cycling time for the feedback collateral system that connects a local group of pyramidal neurons [26, 27, 37, 38]. Consistent with this, the associational projection contains a high proportion of di-synaptic motifs and displays a high efficacy of synaptic signaling at these connections [28]. The spacing (5ms) between spikes is not an unlikely value for the time needed for a pyramidal cell to generate a spike in response to input and then activate another cell. Stereotypy implies that signals are transmitted as packets with information encoded by a specific collection of active CA3 cells rather than as spike patterns. Sparsification in the DG followed by amplification in CA3 via massively branching collaterals would likely present difficulties in obtaining the same spatial pattern of responding cells to multiple applications of the same cortical input. The direct path, which bypasses the DG, provides a presumably more reliable transfer of the cortical signal to CA3 and could bias the response to indirect path activation (by the same cortical input) in favor of a single population of pyramidal neurons. SPWs on the other hand are mostly triggered by stochastic release from the mossy fibers and thus necessarily involve different cells from one wave to the next. The spacing mechanism proposed here to reflect cycling time will nonetheless produce a reasonably constant number of cells to include in successive SPWs.

Despite the slow conduction of the mossy fibers, there were no reliable differences in the time to peak for recurrent responses recorded at three sites along the proximo-distal axis of CA3. The associational fibers, in addition to their dense local ramifications, have rapidly conducting branches that travel across the full extent of CA3. The latter might be thought of as a serial circuit that is parallel to, but faster than, the mossy fibers. We suggest that the long associational projections from those segments of CA3 first activated by the DG „catch up‟ with conduction along the unmyelinated mossy fibers as the latter travel along the proximo-distal axis and shorten the delay to the onset of the recurrent response. The net result of these features will be a near synchronous output from the full extent of CA3 to the different segments of CA1. The direct LPP projection to CA3 adds a type of parallel circuit to the two serial circuits just noted. As discussed, this input could influence which CA3 cells will respond to the collection of cortical inputs that are activated by a particular environmental signal.

The observation that the theta frequency activity was processed through the entire network with minimal distortion accords with the idea that the theta rhythm serves as a carrier wave for the cortical telencephalon [10]. The circuit rejected gamma frequency trains as expected from earlier work showing that the LPP synapses with the DG perform low pass filtering of incoming signals [39]. However, brief gamma bursts triggered a reliable CA1 output when the bursts were separated by 200ms (theta-gamma). These results point to a curious feature of the response to repetitive input: the output of the circuit to theta or theta-gamma activation of the LPP was not greatly different than the input signal. Thus, despite the capacity of the individual links and nodes to perform a diverse array of signal transformations, the circuit, in the stripped down state found in a slice, simply transmits a largely unchanged copy of an acceptable input to downstream targets. This suggests that the many and disparate constituents of the system are in some senses balanced so that net signal transformations are held to a minimum. If so, then the numerically minor afferents from the septum, thalamus, and brain stem [14] could disturb this default mode and produce profound operational changes with relatively discrete effects on select elements. Adjustments to frequency dependent CA3 loops provide one route whereby the coincidence detection and associated spike integration windows can be adjusted during repetitive input. Field CA2, which is innervated by select lower brain areas, has excitatory and inhibitory feed-forward and feedback loops [40–42] that likely perform additional signal modifications [43, 44]. The circuit would thus have multiple modes of operation to be engaged in a situation dependent manner. The throughput mode described here may be employed in familiar environments where active encoding is not required. The default signal would serve to confirm that the contents of the environment or events are as they should be. Activation of previously encoded place fields in a familiar environment, which is disrupted by suppression of the CA3 associational system [45], is an example of such a scenario. Other circumstances would in this model lead to activation of ascending inputs and adjustments so that output is a considerably transformed copy of the input.

The multi-mode arguments in the above hypothesis would hold for many types of complex systems and accordingly do not account for the unusual characteristics of hippocampus. The latter are presumably substrates for specialized operations of a type that cannot be performed by more conventional cortical architectures. SPWs are a striking example of a physiological outcome generated by singular elements. The frequency of the waves depends on stochastic release by the exotic mossy fiber terminals while production and propagation are mediated by the exceptionally dense feedback system that is a defining feature of CA3 [37, 38]. The functional significance of the waves has been a topic of considerable speculation and experimental work [10, 46, 47] but it is reasonable to assume that minutes long periods of repetitive stimulation by large numbers of hippocampal neurons will have important consequences for downstream targets, especially when they are repeated throughout the day-night cycle. Other work has shown that activation of the CA3 associational projections can trigger reverberating activity lasting for extraordinary periods [9]. The system thus approximates recurrent networks of the type hypothesized by Hebb (the 2^nd^ postulate) and others to deal with the problem of associating cues separated by much longer intervals than those required for operant leaning. Depressing the recurrent activity by unilateral silencing of small segments of CA3 pyramidal neurons prevented mice from acquiring the sequence in which cues were sampled in an episodic learning paradigm without affecting encoding of the identities and locations of the cues [9]. The circuit could thus add the critical temporal ordering element to an episodic memory. However, activation of reverberating activity in the above experiments occurred in less than 40% of trials and there are no results suggesting that it can be initiated by cortical input. It may well be the case that minority inputs from the lower brain are needed to shift the circuit into a mode in which cues elicit prolonged firing and sequences are encoded.

## Methods

### Animals

All studies used male C57/BL6 mice (Charles River) from 2-4 months of age. Animals were group housed (5 per cage) with access to food and water ad libitum and were on a 12-h light/dark cycle, with lights on at 6:30AM. Experiments were conducted in accordance with the Institutional Animal Care and Use Committee at the University of California, Irvine and the National Institute of Health Guidelines for the Care and Use of Laboratory Animals. For all electrophysiology studies, mice were anesthetized with isoflurane and euthanized by decapitation.

### Electrophysiological recordings

Hippocampal slices were prepared as previously described [9, 39]. For all electrophysiological hippocampal studies, recordings were digitized at 20kHz using an AC amplifier (A-M Systems, Model 1700) and collected using NacGather 2.0 (Theta burst Corp.).

*Signal throughput CA1:* A stimulating electrode was placed in the dentate gyrus (DG) outer molecular layer towards the apex of the two granule cell blades targeting the direct and indirect LPP projections while two recording pipettes were positioned, the first in CA1c str. radiatum and the second in str. pyramidale of the same subfield (**Fig 1a**). Responses (fEPSP, single units) to single pulse and repetitive stimulation trains (i.e., 5hz, 50Hz) and patterns (i.e., theta-gamma) were recorded. A subset of experiments used surgical cuts to test the relative contributions of the direct and indirect LPP paths to the CA1 response (i.e., fEPSP, spiking), while the LPP-evoked spiking to single-pulse stimulation was investigated in field CA3a.

*MF-CA3 responses:* to directly activate MF projections, a stimulating electrode was positioned within the hilus proximal to the granule cell layer. MF-evoked fEPSP were recorded from the dendritic field and cell body layer of CA3c. Single units were readily detected in the cell body layer response. Stimulation intensity was set to evoke a modest mono-synaptic fEPSP in the PC layer. Responses across the proximo-distal axis of CA3 were recorded in a second set of experiments where pipettes were positioned on the apical edge of the PC layer in CA3c, CA3b and CA3a (**Fig 3a**). See **Suppl. Materials** for a more detailed description of electrophysiological recordings.

### Analysis of data

All recordings were analysed off line. The properties of the fEPSP waveform were analyzed using NacShow 2.0 (Theta Burst Corp) while a custom code (Python version 3.8) was used to analyze spontaneous hippocampal activity (i.e., single units, SPWs) and evoked (single-pulse and repetitive stimulation) single unit activity (see **Suppl. Material** for more details).

