## Supplemental Material for "Input / Output Relationships for the Primary Hippocampal Circuit"

Supplemental Methods.

*Animals*

All studies used male C57/BL6 mice (Charles River) from 2-4 months of age. Animals were group housed (5 per cage) with access to food and water ad libitum and were on a 12-h light/dark cycle, with lights on at 6:30AM. Experiments were conducted in accordance with the Institutional Animal Care and Use Committee at the University of California, Irvine and the National Institute of Health Guidelines for the Care and Use of Laboratory Animals. For all electrophysiology studies, mice were anesthetized with isoflurane and euthanized by decapitation.

*Electrophysiological recordings*

Hippocampal slices were prepared as previously described (Cox et al., 2019; Quintanilla et al., 2022). Experiments were initiated from 8-10AM. Upon removal from the cranium, brains were placed in ice cold, oxygenated (95% O_2_/ 5% CO_2_) high Mg^2+^, artificial cerebrospinal fluid (HM-aCSF) containing (in mM): 87 NaCl, 26 NaHCO_3_, 25 glucose, 75 sucrose, 2.5 KCl, 1.25 NaH_2_PO_4_, 0.5 CaCl_2_, 7 MgCl_2_, (320-335 mOsm). Horizontal sections (400μm) were cut using a Leica Vibrotome (model VT1000s, Deer Park, IL, USA) in ice cold (4^o^C) HM-aCSF and rapidly transferred to an interface recording chamber containing a constant perfusion (60-70 ml/hr) of oxygenated (95% O_2_/ 5% CO_2_) aCSF containing (in mM): 124 NaCl, 3 KCl, 1.25 KH_2_PO_4_, 1.5 MgSO_4_, 26 NaHCO_3_, 2.5 CaCl_2_, and 10 glucose (300-310 mOsm, pH 7.4, 31±1°C). Recordings began 1-1.5hrs later. For all extracellular hippocampal studies, recordings were digitized at 20kHz using an AC amplifier (A-M Systems, Model 1700) and collected using NacGather 2.0 (Theta burst Corp.).

*Signal throughput CA1:* A stimulating electrode was placed in the dentate gyrus (DG) outer molecular layer towards the apex of the two granule cell blades targeting the direct and indirect LPP projections while two recording pipettes were positioned, the first in CA1c str. radiatum and the second in str. pyramidale of the same subfield (**Fig 1a**). Single pulse stimulation produced a complex two part field excitatory postsynaptic potential (fEPSP) and stimulation intensity was set such that single unit was reliably observed on the second component in the CA1 pyramidal cell (PC) layer response. A 20-minute baseline period of single pulse stimulation (0.3Hz) was recorded, with spontaneous activity (i.e., single units, sharp waves [SPW]) additionally collected (10-seconds) after each stimulation pulse. To test the effect that repetitive LPP stimulation had upon CA1 spike output, brief trains were delivered across a range of frequencies (i.e., theta [5Hz], gamma [50Hz]) or patterns (theta-gamma; 5 bursts). Stimulation trains were delivered in a random order and separated by a minimum of 10 minutes. To confirm stimulating electrode placement fEPSPs in stratum lacunosum of field CA3 were recorded in response to paired pulse stimulation (40ms interval): only slice displaying robust paired pulse facilitation, indicating that the activated fibers belong to the LPP rather than MPP (Barrionuevo; Quintanilla; Others), were included in the analysis.

*Contribution of direct & indirect paths:* to test the relative contributions of the direct and indirect paths to CA1 spike output surgical cuts were made to the hippocampal slice to sever the mossy-fiber (MF)-CA3 projection and the LPP-CA3 projection respectively (**Suppl. Fig 1b**). Surgical cuts were made immediately prior to slices being placed on the interface chamber to recover (for 1-1.5-hrs) following slice preparation. CA1 responses to baseline single pulse and repetitive stimulation (i.e., theta gamma and theta-gamma) were conducted as described above. Following each experiment, MF-CA3 cuts were confirmed by delivering a 20Hz (10 pulse) train via a stimulating electrode re-positioned in the MF projections and recording responses from a pipette placed in the PC layer of CA3b (**Suppl. Fig 1b**). LPP-CA3 cuts were confirmed by repositioning one of the recording pipettes to the distal apical dendrites (i.e., stratum lacunosum moleculare) of CA3, and delivering paired-pulse stimulation (40ms interval. **Suppl. Fig 1c**).

*MF-CA3 responses:* to directly activate MF projections, a stimulating electrode was positioned within the hilus proximal to the granule cell layer and fEPSPs recorded from field CA3c via two recording pipettes, the first in str. radiatum and the second on the apical edge of the PC layer. Stimulation intensity was set to evoke a modest mono-synaptic fEPSP in the PC layer. A second set of experiments recorded responses across the proximo-distal axis of CA3. As such recording pipettes were positioned on the apical edge of the PC layer in CA3c, CA3b and CA3a (**Fig 3a**).

*LPP-CA3 responses:* a stimulating electrode was positioned in the dentate gyrus (DG) outer molecular layer (as above) and a single recording pipette positioned within the PC layer of CA3a. The stimulation intensity was set such that a modest fEPSP (0.5-1mV) was evoked that reliably contained single units. A 20-minute period of single pulse stimulation (0.3Hz) was recorded, with spontaneous activity (i.e., single units, sharp waves [SPW]) additionally collected (10-seconds) after each stimulation pulse.

*Analysis of data*

All recordings were analysed off line. The properties of the fEPSP waveform were analysed using NacShow 2.0 (Theta Burst Corp), while a custom code (Python version 3.8) was used to analyze spontaneous hippocampal activity (i.e., single units, SPWs) and evoked single unit activity.

*fEPSP analysis:* LPP-evoked fEPSPs recorded from the apical dendrites (i.e., str. radiatum) of CA1c were analyzed with regard to peak amplitude (total), the initial slope (20-80%) of the rising phase of the first and secondary fEPSP components, the area and half width of the waveform. An ensemble average fEPSP was generated (20 events) and used to calculate the fEPSP decay τ and latency to onset. The decay τ was described using the mono-exponential equation *Y(t)=A*exp(-1/t)*. Responses recorded from the cell body layer (i.e., str. pyramidale) were analyzed with regard to peak amplitude and area. Total values as well as those for the initial and secondary components of the waveform were measured.

*LPP-evoked single units:* LPP- and MF-evoked spikes recorded from CA1 or CA3 were analyzed offline using a custom-written computer code created with Python 3.8. Briefly, extracellular recordings (10 second sweeps) were fed through a band pass filter (300-5000Hz). Spikes were detected using a combined amplitude threshold (-45μV) and rate of rise threshold (100μV/ms). The spike output generated by single-pulse LPP activation was analyzed across the 50ms period following the stimulation artifact. For each slice 30 consecutive responses to single pulse stimulation were analyzed. Responses were analyzed with regard to number of spikes, inter-spike interval (ISI), latency to 1^st^ spike, spike amplitude and instantaneous frequency (i.e., 1/ISI) of spike output. The standard deviation of the 1^st^ spike latency was used to describe the “jitter” within each slice. To confirm the measured spikes were driven by LPP activation we calculated the probability of such an output occurring spontaneously after any given spontaneous spike. For each slice we took the mean ± SD of the spike output (number and frequency) along with latency to first spike and measured the probability of such a pattern occurring after any single spontaneous spike across the 10 seconds of each sweep (30 consecutive. **Suppl. Table 2 & 4**).

*Analysis of repetitive stimulation*: the CA1 spike output to brief repetitive LPP stimulation at different frequencies (i.e., 5Hz, 50Hz) and patterns (i.e., theta gamma) was analyzed using the same custom code described above. LPP-evoked spikes were analyzed for each pulse with regard to number of spikes, inter-spike interval (ISI), latency to 1^st^ spike (and associated “jitter’), spike amplitude and instantaneous frequency (i.e., 1/ISI) of spike output. For all 5Hz stimulation, spiking was analyzed for 50ms following the stimulation artifact associated with each pulse. Responses to LPP stimulation at 50Hz or with theta-gamma bursts, spiking was assessed i) across the 20ms interval between gamma frequency pulses and ii) over 50ms periods following the initial pulse in the train (i.e., divide into four consecutive 50ms epochs) or burst. Assessing LPP-evoked spiking over such 50ms periods enabled quantitative comparison with single-pulse output (above) as well as with the response to 5Hz stimulation.

Supplemental Figures

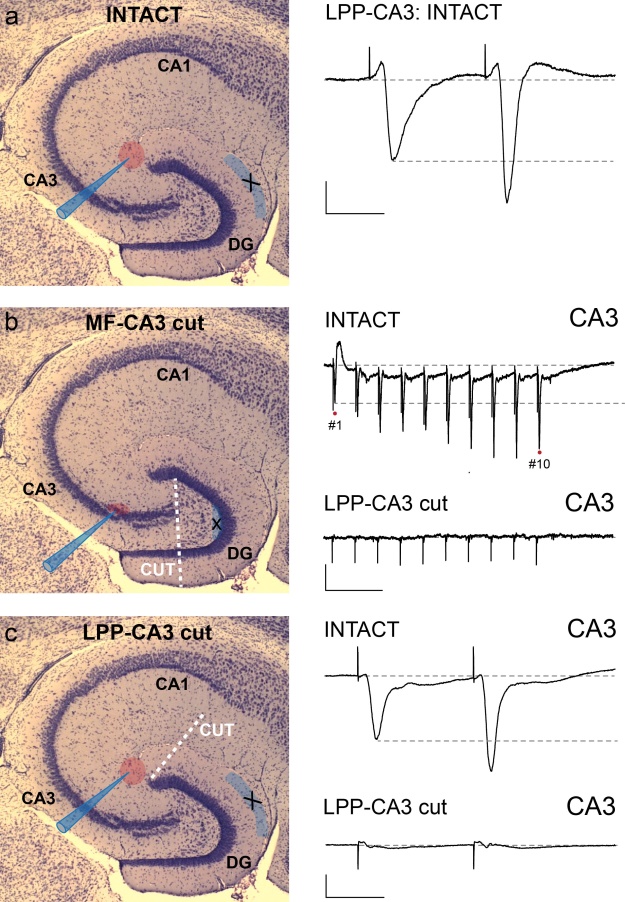

**Suppl. Figure 1. Electrode placements for the different circuit configurations.** (Right) Nissl stains of hippocampal slices depicting the stimulating electrode placement and position of recording pipettes in the intact slice (**a**), slices where the MF-CA3 projection was cut (i.e., direct LPP only. **b**), and slices where the LPP-CA3 was cut (i.e., indirect LPP only. **c**). Example responses to confirm placement and completeness of cuts are illustrated for each (right). Scale bars: **a**: y=0.5mV, x=20ms; **b**: y=0.5mV, x=100ms; **c**: y=1mV, x=20ms.

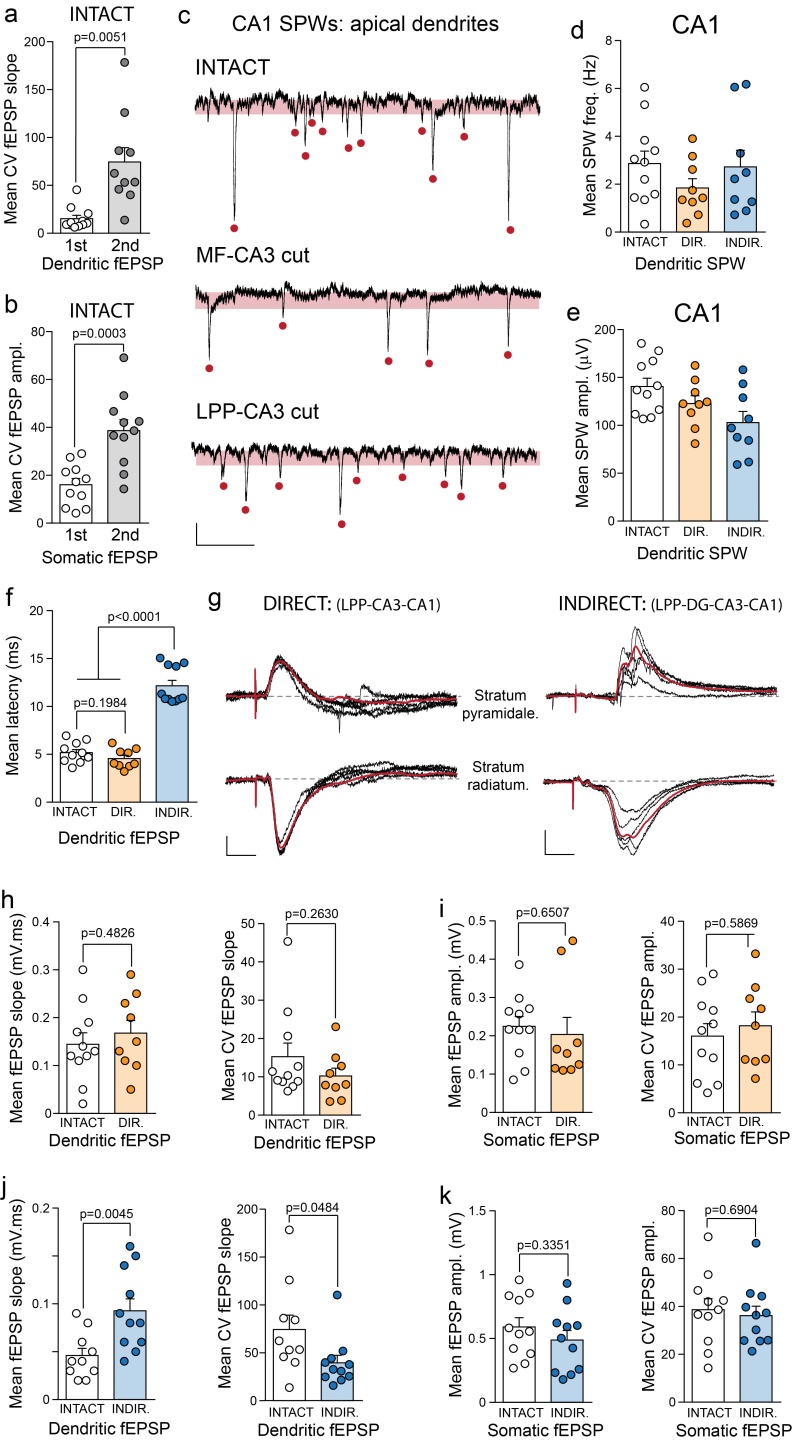

**Suppl. Figure 2. The direct and indirect LPP contribute differentially to a two-part fEPSP in CA1.** Graphs summarizing the mean CV of (**a**) the slope of the two components of the dendritic fEPSP and (**b**) the peak amplitude of the two components recorded from the PC layer. **c.** Representative SPWs recorded from the apical dendrites of intact slices (top) and those lacking the indirect LPP input (i.e., MF-cut; middle) and the direct LPP input (i.e., LPP-CA3 cut; bottom). Bar graphs summarizing the mean frequency (**d**) and amplitude (**e**) of SPWs recorded from the CA1 apical dendrites in each slice configuration. The amplitude of SPWs in slices lacking the direct LPP input was modestly yet significantly decreased *cf* the intact circuit (p=0.0163 unpaired t test). Individual events are depicted (red circles). **f**. Graph summarizing the mean latency to the initial onset of the dendritic fEPSPs recorded from each slice configuration. **g**. Representative LPP-evoked fEPSPs recorded from the PC layer (top) and apical dendrites (bottom) of CA1 in slices containing only the direct (left) or indirect LPP (right) input. The ensemble average fEPSP for each is depicted in red (scale bars: y=0.1 and 0.25mV, x=10ms). Bar graphs comparing the mean slope (left) and associated CV (right) of the dendritic fEPSP (**h, j**) and mean amplitude (left) and associated CV (right) of the somatic fEPSP (**I, k**) recorded from CA1 of slices containing the direct (orange: **h, i**) or the indirect (blue: **j, k**) LPP path only, with the initial component of the response in the intact slice.

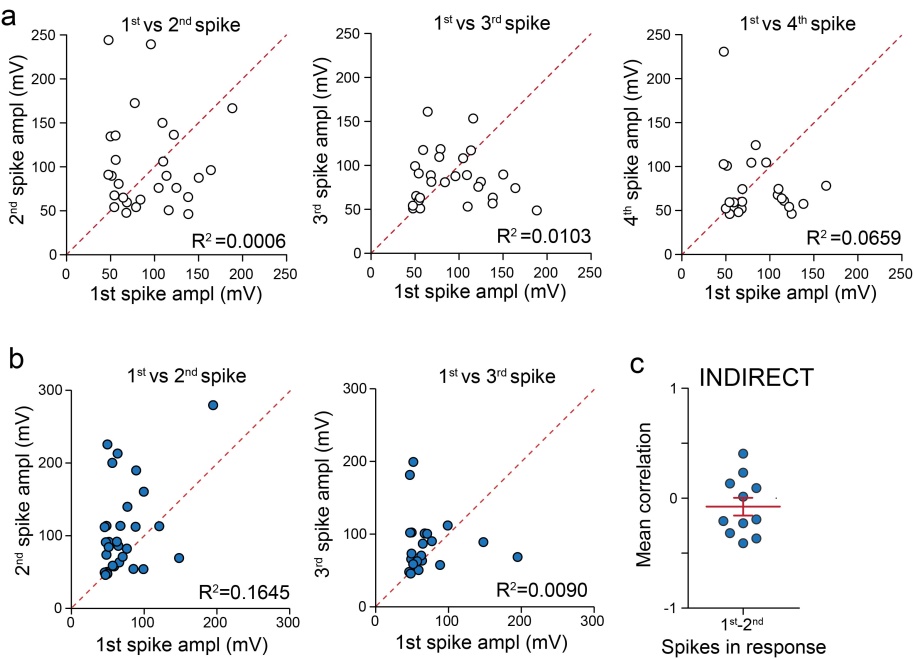

**Suppl. Figure 3. Activation of the circuit in the absence of the direct LPP path produces a CA1 spike output similar to the intact circuit.** Scatter plots with unity lines (red dashed) summarizing (**a**) the correlation of the 1^st^ spike amplitude with the 2^nd^, 3^rd^ and 4^th^ spikes (across 30 successive trials) spikes in the intact circuit (open circles) and (**b**) the correlation between the amplitude of the 1^st^ spike with the 2^nd^ and 3^rd^ spikes in a representative slice containing only the indirect input (blue circles). Graph summarizing the mean correlation between the amplitude of the 1^st^ and 2^nd^ spikes recorded for all those slices lacking the direct LPP input i.e., indirect path only (**c**).

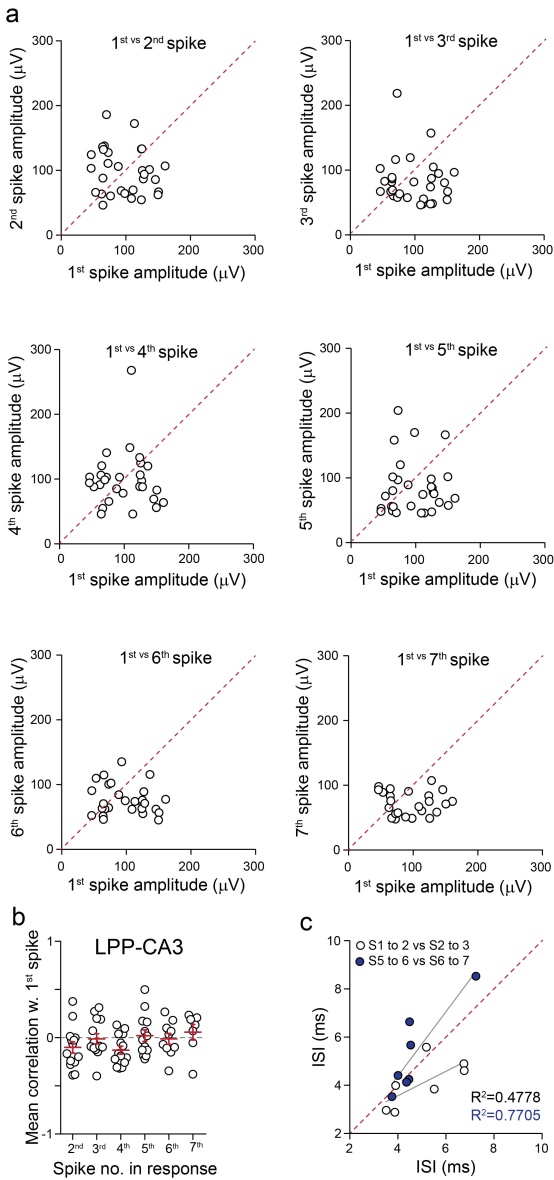

**Suppl. Figure 4. LPP activation produces a stereotyped CA3 spike output from a small population of cells. a.** Scatter plot with unity lines (red dashed) summarizing the correlation between the 1^st^ spike amplitude with all subsequent spikes (up to 7) within the response from 30 successive trials recorded from a representative slice. **b**. Graph summarizing the mean correlation between the 1^st^ spike amplitude with all subsequent spikes (up to 7) from all slices. **c.** Scatter plot with unity lines (red dashed) summarizing the correlation between the mean ISI values between the 1^st^-2^nd^ and the 2^nd^-3^rd^ spikes (open circles) as well as the 5^th^-6^th^ and the 6^th^-7^th^ spikes (blue circles) recorded from CA3 following single-pulse LPP stimulation.

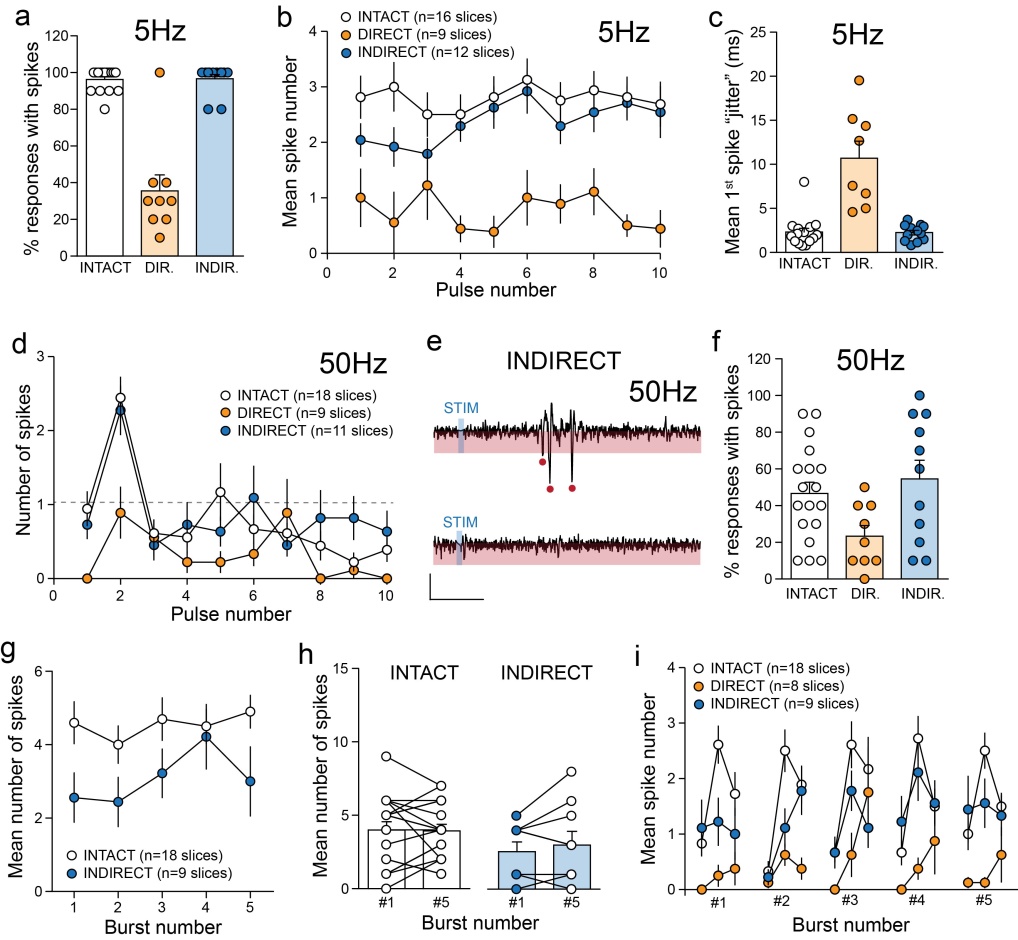

**Suppl. Figure 5. Signal throughput following repetitive stimulation is dependent upon the frequency and pattern of LPP activation. a.** Graph summarizing the proportion (%) of CA1 responses containing single units during repetitive LPP stimulation at 5Hz for each circuit configuration (i.e., intact, direct and indirect paths). **b**. Graph showing the mean number of spikes elicited for each pulse in a 5Hz (10 pulse) train in the intact circuit and with only the direct or indirect path present. **c**. Graph illustrating the mean “jitter” associated with the latency to first spike across the 10 successive pulses for slice in the different circuit configurations. **d**. Graph showing the mean number of spikes elicited for each pulse in a 50Hz (10 pulse) train in the intact circuit and with only the direct or indirect path present. **e**. Representative filtered baseline CA1 response (top) and 10^th^ pulse (bottom) of a 50Hz (10 pulse) stimulation train evoked by the indirect path only (scale bars: y=50μV; x=10ms). **f**. Graph summarizing the proportion (%) of CA1 responses containing single units during repetitive LPP stimulation at 50Hz for each circuit configuration (i.e., intact, direct and indirect paths). **g**. Graph illustrating the mean CA1 spike number per burst elicited by theta-gamma activation of the LPP in the intact circuit (open circles) and with the indirect path only (blue circles). **h**. Bar graphs summarizing the mean number of spikes evoked by the 1^st^ and 5th bursts in a theta-gamma train in slices containing the intact circuit (open circles) and only the indirect LPP input (blue circles). **i**. Graph showing the mean number of CA1 spikes elicited by each pulse of a theta-gamma (5 burst) train in the intact circuit and with only the direct or indirect path present.

A. DENDRITIC response

|  | INTACT circuit  (n=11 slices) | DIRECT Circuit  (n=9 slices) | INDIRECT Circuit  (n=11 slices) |
| --- | --- | --- | --- |
| fEPSP ampl. (mV) TOTAL | 0.60 ± 0.08 | 0.42 ± 0.05 | 0.58 ± 0.05 |
| fEPSP slope (mV.ms) INITIAL | 0.14 ± 0.02 | 0.17 ± 0.03 | - |
| fEPSP slope (mV.ms) SECOND | 0.05 ± 0.01 | - | 0.09 ± 0.01** |
| Area (mV.ms) | 9.4 ± 1.6 | 3.7 ± 0.5** | 9.5 ± 0.8 |
| Half width (ms) | 11.7 ± 1.0 | 7.3 ± 0.9** | 14.2 ± 04* |
| Decay τ (ms) | 7.9 ± 1.6 | 5.4 ± 0.7 | 8.9 ± 0.7 |
| Latency to onset (ms) | 5.2 ± 0.3 | 4.6 ± 0.3 | 12.2 ± 0.6*** |

B. SOMATIC response

|  | INTACT circuit  (n=11 slices) | DIRECT Circuit  (n=9 slices) | INDIRECT Circuit  (n=11 slices) |
| --- | --- | --- | --- |
| fEPSP ampl. (mV) TOTAL | 0.60 ± 0.07 | 0.22 ± 0.04 | 0.49 ± 0.08 |
| fEPSP ampl. (mV) INITIAL | 0.24 ± 0.02 | - | - |
| fEPSP ampl. (mV) SECOND | 0.59 ± 0.07 | - | - |
| Area (mV.ms) TOTAL | 10.9 ± 1.3 | 2.2 ± 0.5 | 7.5 ± 1.3 |
| Area (mV.ms) INITIAL | 1.4 ± 0.2 | - | - |
| Area (mV.ms) SECOND | 9.5 ± 1.3 | - | - |

**Supplemental Table 1.** Properties of dendritic (A) and somatic (B) LPP-evoked fEPSPs recorded from slices containing the intact hippocampal circuit as well as those lacking the indirect or direct components.

| CA1 spike properties | INTACT circuit  (n=20 slices) | DIRECT Circuit  (n=9 slices) | INDIRECT Circuit  (n=13 slices) |
| --- | --- | --- | --- |
| # of spikes | 2.7 ± 0.3 | 0.6 ± 0.3 | 2.1 ± 0.3 |
| Max # of spikes | 5.0 ± 0.3 | 3.1 ± 0.6 | 4.2 ± 0.4 |
| 1^st^ spike latency (ms) | 19.6 ± 1.0 | 25.4 ± 2.8* | 20.4 ± 0.9 |
| “jitter” 1^st^ spike latency (ms) | 4.2 ± 0.4 | 10.4 ± 2.0*** | 3.3 ± 0.5 |
| 1^st^ spike amplitude (μV) | 95.8 ± 8.3 | 80.4 ± 10.1 | 90.7 ± 9.7 |
| % responses w. spikes | 92.2 ± 2.3 | 23.7 ± 7.5*** | 89.8 ± 3.6 |
| % responses w. ≥2 spikes | 76.0 ± 5.1 | - | 67.3 ± 7.5 |
| Instantaneous frequency (Hz) | 255.9 ± 11.9 | - | 259.5 ± 14.5 |
| Prob. spont. spike pattern ALL | 0.81 ± 0.3 | - | 0.80 ± 0.4 |
| Prob. spont. spike pattern in SPW | 0.11 ± 0.03 | - | 0.05 ± 0.03 |
| Prob. spont. spike pattern o/s SPW | 0.70 ± 0.3 | - | 0.75 ± 0.3 |

**Supplemental Table 2.** Properties of LPP-evoked CA1 spiking recorded from slices containing the intact hippocampal circuit as well as those lacking the indirect or direct components.

|  | SOMATIC response | | DENDRITIC response |
| --- | --- | --- | --- |
| fEPSP properties | Mono-synaptic MF response  (n=11 slices) | Di-synaptic MF response  (n=6 slices) | Di-synaptic MF response  (n=6 slices) |
| Amplitude (mV) | 0.5 ± 0.2 | 2.3 ± 0.2 | 3.3 ± 0.4 |
| Slope (mV.ms) | - | 1.3 ± 0.2 | 1.5 ± 0.3 |
| Area (mV.ms) | - | 21.5 ± 2.4 | 39.0 ± 4.6 |
| Half width (ms) | - | 8.3 ± 0.7 | 7.8 ± 0.6 |
| Decay τ (ms) | - | 6.1 ± 0.6 | 11.5 ± 0.8 |
| Latency to onset (ms) | 3.2 ± 0.2 | - | - |
| Latency to peak (ms) | 4.4 ± 0.2 | 8.5 ± 0.6 | 8.0 ± 0.5 |

**Supplemental Table 3.** Properties of mono- and di-synaptic MF-evoked fEPSPs recorded from cell body (somatic) and dendritic layers of field CA3a. Note that mono-synaptic MF-evoked responses were only recorded from the cell body layer of this subfield.

| CA3 spike properties | LPP-CA3  (n=14 slices) | MF-CA3  (n=6 slices) |
| --- | --- | --- |
| Mean # of spikes | 6.4 ± 0.5 | 3.7 ± 1.0* |
| Mean max # of spikes | 9.9 ± 0.7 | 6.8 ± 1.7 |
| 1^st^ spike latency (ms) | 8.3 ± 0.4 | 4.9 ± 0.4*** |
| “jitter” 1^st^ spike latency (ms) | 3.3 ± 0.3 | 0.9 ± 0.2*** |
| 1^st^ spike amplitude (μV) | 89.7 ± 5.1 | 236.7 ± 41.7*** |
| % responses w. spikes | 100 ± 0.0 | 100 ± 0.0 |
| % responses w. ≥2 spikes | 99.5 ± 0.5 | 91.7 ± 4.1* |
| Instantaneous frequency (Hz) | 273 ± 5.3 | 328 ± 24.0** |

**Supplemental Table 4.** Properties of LPP- and MF-evoked spiking recorded from CA3a.
